# Wavefunction Patterns the Embryo?

**DOI:** 10.1101/2025.08.22.671254

**Authors:** Irfan Lone

## Abstract

The problem of cell fate determination by morphogen gradients in the embryonic development of many multicellular organisms has been a long-standing and important one in developmental biophysics and dynamics. The first mathematical model was proposed by Francis Crick over 50 years ago by postulating a reaction-diffusion based mechanism underlying the whole process. The first real world morphogen, named Bicoid (Bcd), was identified by molecular biologists in late 1980s in the embryo of fruit fly *Drosophila melanogaster*. Subsequently, Crick’s classical random walk based model was used by biophysicists to explain the formation of Bicoid and other morphogen gradients. Very recently Fluorescence correlation spectroscopy (FCS) studies have revealed multiple modes of Bcd transport at different spatial and temporal locations across the embryo of Drosophila melanogaster. It has been be shown that these observations are best fitted by a model based on quantum mechanics. In such a model it is hypothesized that the transitory quantum coherences in collaboration with unitary noise are responsible for the observed dynamics and relaxation to a non-equilibrium steady-state of the Bcd morphogen gradient. In this paper, in addition to further clarifying the mathematical details underlying the quantum-classical model, we use the said model to explain the observed Bcd interpretation time by its primary target gene Hunchback (*hb*).

---

A morphogen gradient is defined as a concentration field of a molecule that acts as a dose-dependent regulator of cell differentiation [1–7]. Bicoid (Bcd) gradient is a prototypical morphogen gradient that is formed during the early embryogenesis of fruit fly *Drosophila melanogaster* and is known to play a crucial role in the determination of cell fate and polarity in the organism [8–11]. Despite having been studied for a very long time, the biophysical mechanisms behind the establishment of the Bicoid gradient are still not well understood.

Over 50 years ago, Francis Crick suggested that morphogen gradients could be established through molecular diffusion and degradation by introducing a model in which there is a constant production of morphogen and its spatially uniform degradation throughout the developing system with the consequence of setting up a concentration gradient in three-dimensional space [12]. Mathematically, such a system is described through the use of following reaction-diffusion equation,

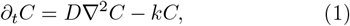

where *D* and *k*, respectively, denote the diffusion and degradation rate of the morphogen, and *C* denotes it concentration [12]. All the subsequent treatments are essentially the extensions of Crick’s ideas [13–29].

In contradistinction to above, recently fluorescence correlation spectroscopy (FCS) and perturbative studies of the Bcd gradient have revealed multiple modes of transport at different spatio-temporal locations across the embryo [30]. In the cytoplasm, for instance, the slow dynamic mode is similar across the embryo while as the fast one shows an increased diffusivity in going from the anterior to the posterior side of the organism. Similarly, the nuclear Bcd shows a significant, though compara-tively small, increase in its fast dynamic mode with the slow component remaining essentially unchanged [30].

These observations, and a few others, can be explained through a quantum-classical treatment of Crick’s model [31]. In such a treatment, the degradation of Bcd is modeled as a unitary noise that is intrinsic and that does not cause any entanglement with the environment, such that the system remains in an essentially pure state during the course of its evolution [31]. Compared to the usual classical random walk models, the coin toss is replaced by a chirality degree of freedom which can take two values denoted |*R*⟩ and |*L*⟩, for right- and left-handed chirality, respectively [32]. A Bcd molecule of chirality |*R*⟩ moves one step to the right at a time while that of chirality |*L*⟩ moves to the left along the lattice (Fig. 1). They can also get degraded, leading to the setting up of a Bcd concentration gradient in three-dimensional space.

**FIG. 1.**
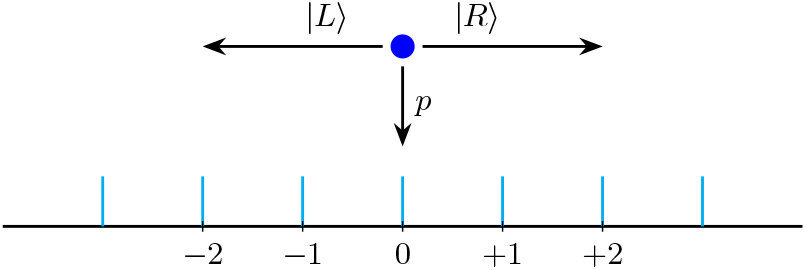
Schematic of the quantum-classical model. A Bicoid molecule (shown by the blue colored dot) can, respectively, make a transition to the left or to the right depending on its chirality state of |*L*⟩ or |*R*⟩ or it can get destroyed as well.

It is known that the time-dependent Schrödinger equation

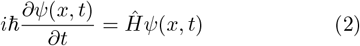

can be applied to the study of chiral molecules [33]. Its general solution is

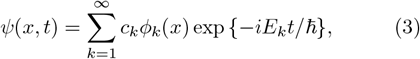

where *c*_*k*_ are complex coefficients, 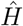 is the Hamiltonian operator, *ϕ*_*k*_(*x*) are the eigenfunctions and *E*_*k*_ are the energy eigenvalues of the stationary states such that

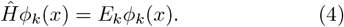

Since all naturally occurring amino acids that go on to constitute the proteins are left-handed, this homochirality ensures that Bcd, just like all other proteins, is a chiral biomolecule [34]. A useful model is to represent a chiral molecule using only the two lowest energy molecular states. The solution of Eq. (2) is then given by

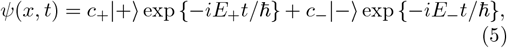

where we have denoted by |+⟩ and |−⟩ the two time-independent Hamiltonian eigenstates [33]. Quantum mechanics, in principle, allows for a coherent superposition of these states that can be used to represent the states |*L*⟩ and |*R*⟩ corresponding to the *L* and *R* enantiomers of a chiral molecule as

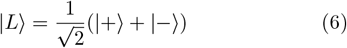

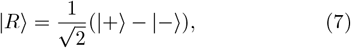

which are therefore called chiral states [33]. Furthermore, the Bcd morphogen inside the embryonic syncytium may be viewed as an ensemble of identical bosons such that the initial state of the system is represented by the superposition given in Eq. (6). Thus, the chirality consists of a single qubit and is a proper representation of the state of the system in this case. This can readily be verified by the application of a Hadamard transformation,

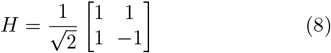

to Eq. (6). Since

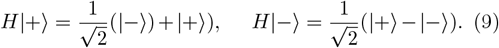

Therefore,

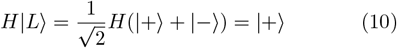

which in view of Eq. (9) again gives back the state |*L*⟩ upon application of the transformation given in Eq. (8). Thus, the choice of initial state represented by Eq. (6) is reasonable one. This choice correctly reproduces the observed statistical features of the Bcd gradient as discussed below.

However, in order to move beyond a trivial discussion of chirality and to explicitly incorporate the spatial degrees of freedom in our treatment we require a more sophisticated approach that, fortunately, also follows in a straightforward manner from the Schrodinger dynamics [35]. In our treatment, the lattice sites, *n* = 0, ±1, ±2, …, correspond to the states of the Bcd particles in the position Hilbert space ℋ_*n*_, which has a support on the space of eigenfunctions ℋ*n*⟩ corresponding to the sites *n* ∈ ℤ, ℤ being a set of integers (Fig. 1). The position Hilbert space ℋ_*n*_ is augmented by a ‘coin’ space ℋ_*c*_ spanned by the two basis states {|*R*⟩, |*L*⟩}. The states of the total system are thus in the Hilbert space

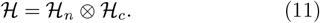

The Bcd molecules move along the integer lattice ℤ in discrete time steps depending on their coin state. The time evolution is therefore expressed by

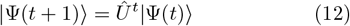

where 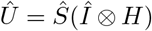 is a unitary operator that defines a step of the walk, 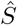 is the conditional shift operator that translates the particle to right or left depending on its chirality state

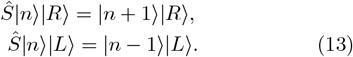

and 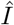 is the spatial identity operator [32]. The quantum state of the Bcd system at a given time *t* is given as

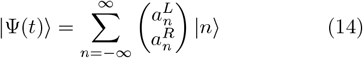

with the amplitude components 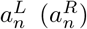 associated to the left (right) chirality states, respectively. For such a pure state the density operator is given by

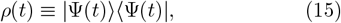

with the normalization

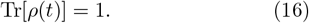

The time evolution of the system can therefore alternatively be described as

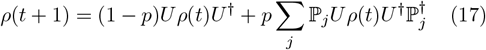

where ℙ_*j*_ is a projection that represents the action of the unitary noise in the form of randomly occurring projective measurements performed on the system and *p* is a probability parameter that quantifies such noise [31]. The probability distribution for finding the walker at site *n* at time *t* is thus calculated by using

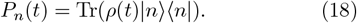

The derivation of the expression for the diffusion coefficient *D* characterizing such a distribution is somewhat cumbersome, albeit straightforward.

Let ℋ_*e*_ denote the Hilbert space of the environment spanned by the orthonormal basis 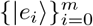, where *m* is the dimension of the environment Hilbert space. The total Hilbert space, including that of the environment, is thus given by

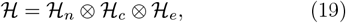

The state of the system is then obtained by tracing out over the environmental degrees of freedom as

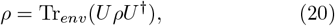

Now one may assume that the initial state of the whole system is given by the tensor product

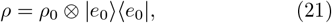

where *ρ*_0_ gives the state of the principal system and |*e*_0_⟩⟨*e*_0_| is the environmental state. Then one can write

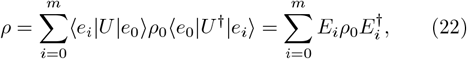

where *E*_*i*_ = ⟨*e*_*i*_|*U*|*e*_0_⟩, *i* = 0, 1, …, *m*, are the so-called Kraus operators. The Kraus operators follow the completeness relation, which arises from the requirement that

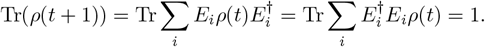

Since above equation is true for all *ρ* then

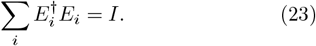

From the definition of Kraus operators in Eq. (22), a step of the walker is given by

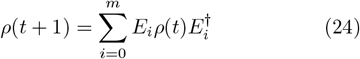

For *t* steps one has

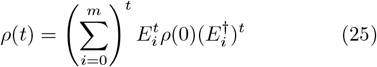

The coin operator, the translation operator and the environment effects are all embedded in *E*_*i*_. Our task is then reduced to finding the appropriate Kraus operators for our system defined in Eq. (22) and then use Eq. (25) in order to obtain the final state *ρ*(*t*).

Let ℛ_*i*_ be the *i*th reaction that affects the system with the corresponding probability *p*_*i*_. Furthermore, let us assume that we have *r* such reaction operators ℛ_*i*_, *i* = 1, …, *r*, where each of them acts on the system with the probability *p*_*i*_. Now, if the environment is in the state |*e*_*i*_⟩ then the operator ℛ_*i*_ acts on the system. Therefore, one can imagine the *r*-dimensional Hilbert space for the environment, spanned by 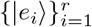, and the following initial state for the environment

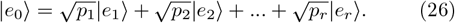

Clearly, the probability of finding the environment in the state |*e*_*i*_⟩ is *p*_*i*_. Therefore one can write the unitary transformation of the whole system (system+environment) in this case as

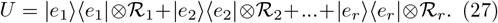

So the Kraus operators will be given by

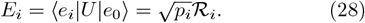

and Eq. (24) gives the density matrix after the first step accordingly as

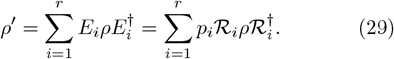

Equation (29) gives *ρ* as a mixture of different evolutions with the corresponding probabilities *p*_*i*_ as expected. Translating this to our setting we can define the initial state of the environment as follows

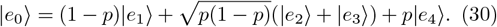

Since the Kraus operators act on the walker Hilbert space, it follows that

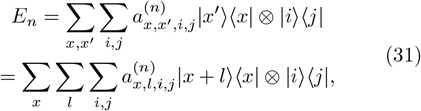

where *x, l* = −∞, …,∞ and *i, j* =, {|−⟩, |+⟩}. Fourier transformation

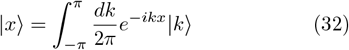

of Eq. (31) turns it into

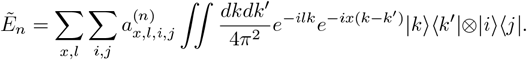

After invoking the homogeneity condition so that the coefficients 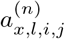 are independent of *x*, above equation takes the form

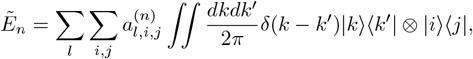

where we have made use of the following orthonormalization relation

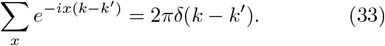

By integrating out the terms involving *k*^*′*^ and changing the order of integration and summation one gets

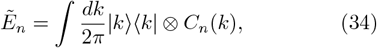

where

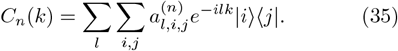

One may, in a similar manner, write the general form of *ρ*_0_ in the *k* space as

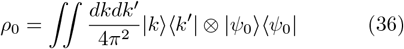

so that Eq. (24) becomes

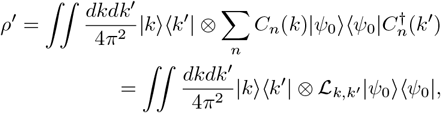

where 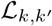 is a superoperator defined as

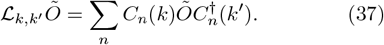

Thus after *t* steps, the state and the probability of finding our Bcd walker at position *x* are, respectively, given by

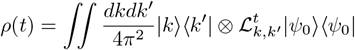

and

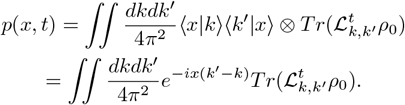

From the completeness relation, Eq. (23), one can thus deduce that

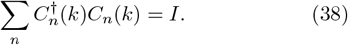

From the above condition it follows that the superoperators ℒ_*k,k*_ are trace preserving, i.e.,

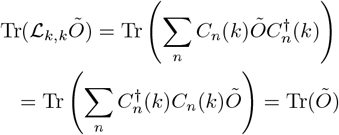

and therefore

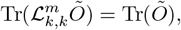

which is true for any arbitrary operator *Õ*. Now the *m*th moment of the probability distribution *p*(*x, t*) is defined by

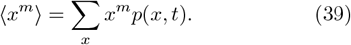

Inserting the above expression for *p*(*x, t*) in Eq. (39) we get

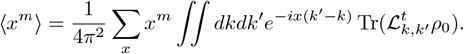

Using the orthonormalization relation (33), one gets for the first and second moments

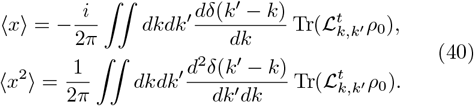

In order to carry out the above integrations one needs the following equations

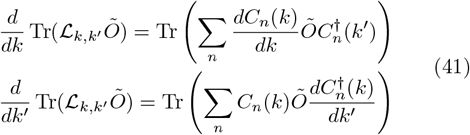

where according to Eq. (35)

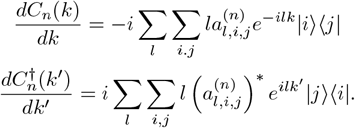

Since the superoperator 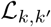 acts on a positive and Hermitian density matrix, one can write Eq. (41) as follows

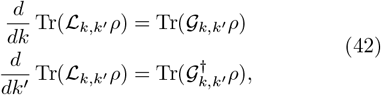

where

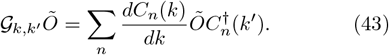

Finally, by carrying out the integrations of Eq. (40), we arrive at the following expressions for the first and second moments

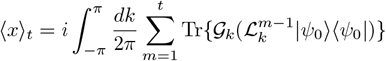

and

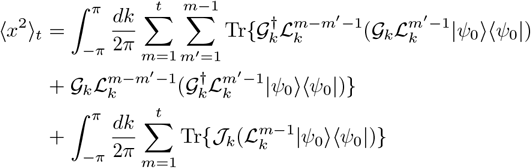

where we have defined

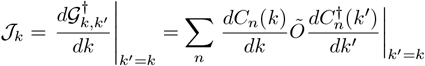

and used 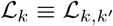 and 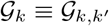 to simplify writing.

Now, following ref. [45], we use the affine map approach in order to find the above superoperators and determine the value of diffusivity as follows. Since the operator ℒ_*k*_ is linear one can represent it as a matrix acting on the space of two-by-two operators. First, any two-by-two matrix can be represented by a four-dimensional column vector as

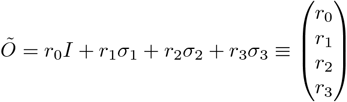

where we have defined *σ*_0_ = *I* and *σ*_*i*_ (*i* = 1, 2, 3) are the usual Pauli matrices. Now, in order to find the above superoperators, we let them operate on an arbitrary two-by-two matrix *Õ* as follows

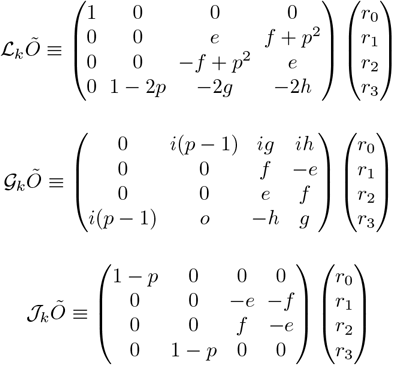

and

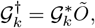

where the last equation is obtained, simply, from the Hermiticity of the Pauli matrices. Here *e, f*, *g*, and *h* are functions of *p* and *k*, defined by

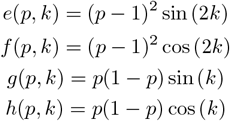

With the above representation in hand, one can calculate the moments given by Eq. (40). It turns out that, in the long time limit, the first moment is independent of time. Our interest here is in finding the diffusion coefficient with the below definition

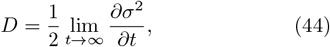

where *σ*^2^ = ⟨*x*^2^⟩−⟨*x*⟩^2^. Since the time independent term does not contribute to the diffusion coefficient *D*, thus we shall focus on finding the second moment in Eq. (40). A somewhat detailed but straightforward calculation shows that the second moment is given by

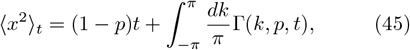

where

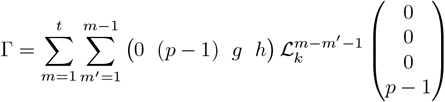

Substituting the result in Eq. (45) into the definition of diffusion coefficient in Eq. (44) and simplifying we have finally

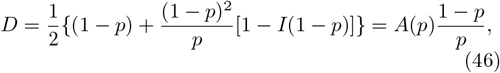

where the coefficient *A* is a function of *p*

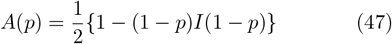

with *A* → 0.5 as *p* → 1, and

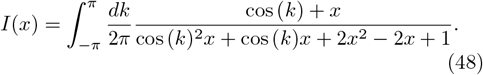

Eq. (46) predicts that the diffusivity of Bcd system is not constant but varies as a function of noise reaching much larger values for very small noise levels (Fig. 2). Such large diffusion coefficients for morphogens are not unusual and values as high as 50 − 80*µm*^2^*s*^*−*1^ have been measured in different developmental contexts [36]. Eq. (46) thus corresponds to the observed fast dynamic mode of the Bcd gradient [30]. Crick, using his source-sink model,

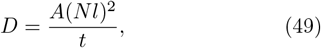

has proposed a general value of *A* = 0.5 for the biochemically more realistic models; where *t* is the time, *N* is the number of cells between the source and sink, and *l* is the length of each cell [12].

**FIG. 2.**
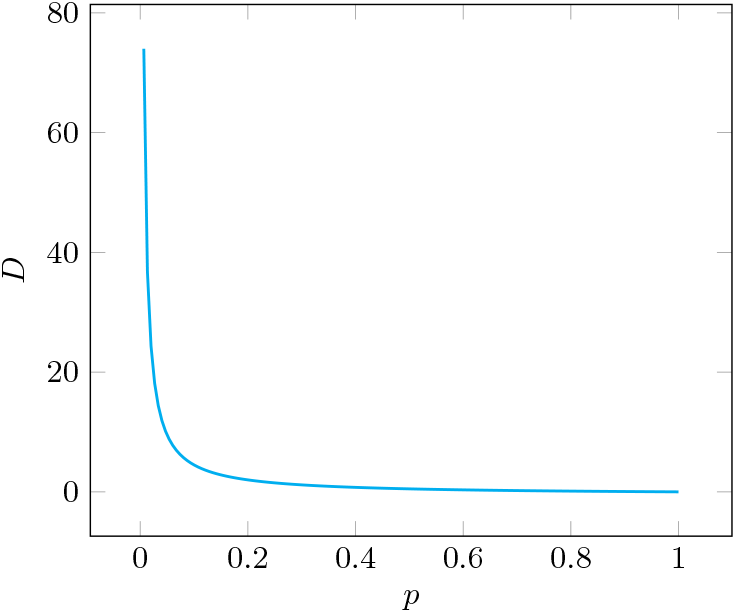
Plot of Eq. (46), as a function of noise, *p*. The magnitude of *D* is large for very small *p*.

The establishment of the Bcd gradient in the 500*µm* long syncytial embryo of Drosophila takes place in about 3 hours [17]. Within about 2 hours the number of nuclei in the embryo increases by three orders of magnitude, from 1 to ∼5000. If one takes the average nuclear diameter, *l* ∼10 *µm* [17], the number of cells along the line between source and sink in the Crick’s model will come out to be about *N* ∼ 50. Using the value of *A* = 0.5, Eq.(49) gives *D* ∼12*µm*^2^*s*^*−*1^, which is the same value as that experimentally observed for the effective diffusivity of Bcd [30]. Thus, the Crick’s source-sink model [12] is able to reproduce the observed dynamics of Bcd system. But this value for diffusivity is too low to explain the expected Bcd interpretation times by its primary target genes like the hunchback (*hb*). The existing theoretical treatments, therefore, predict interpretation times of duration less than a minute and remain unsubstantiated in vivo [37–39]. Experimentally, however, the Bcd protein is known to bind to chromatin sites in a highly transient manner, with specific binding events lasting on the order of a second, in all portions of the embryo [40]. This demands interpretation times of duration less than a second. The quantum-classical model, on the other hand, readily explains such short Bcd interpretation times.

Before turning to the problem of Bcd interpretation and its solution in terms of a quantum-classical picture, we would like to make the observation that as per Eq. (46), the dominant contribution to the diffusion coefficient depends inversely on noise *p*. This dependence is due to the persistence of quantum correlations in the system. In the limit of *p* → 1 and *A* → 0.5, Eq. (46) gives way to

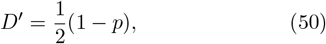

which, in light of Eq. (1), describes a genuine diffusive process with the diffusion equation given by

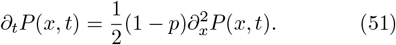

As per Eq. (50), the value of diffusivity (*D*^*′*^) is very small with a maximum at *p* = 0 for a totally classical walk (Fig. 3). Eq.(50) thus corresponds to the observed slow dynamic mode of the Bcd gradient [30].

**FIG. 3.**
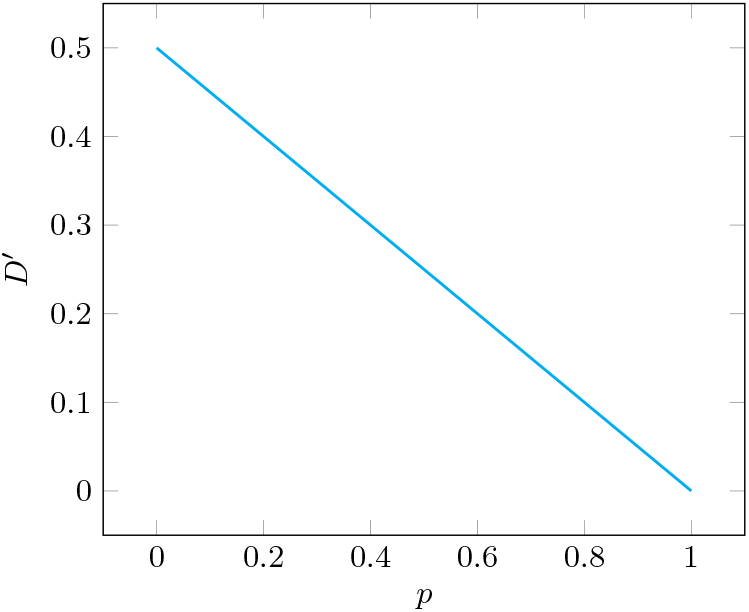
Slow dynamic mode of Bcd, Eq. (50), as a function of noise, *p*. There is no significant variation in *D*^*′*^.

Lastly, it has been shown that the quantitative relationship between the statistics of *hb* expression levels in the embryo is consistent with a model in which the dominant source of noise are the diffusive fluctuations associated with the random arrival of *Bcd* transcription factor molecules at their target sites on the DNA [41]. This noise dependence is quantitatively expressed by the Berg-Purcell limit,

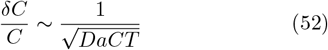

in which *a* denotes the size of the *Bcd* receptor, *T* is the duration of signal integration, *C*(*x, t*) is the local *Bcd* concentration and *D* is the diffusion rate [42]. It is quite conceivable based on Eq. (46) that a significant fraction of Bcd molecules can move much faster given the very low observed degradation rates in the system *k ∼* 10^*−*4^*s*^*−*1^ [43]. This is because the system can vary its diffusivity, depending upon its requirement, by modulating noise [31]. Therefore, substituting Eq. (46) in Eq. (52) one obtains

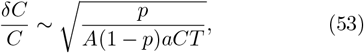

Equation (53) is a modified Berg-Purcell limit for the *Bcd* gradient. For the numerical values of noise *p* comparable in magnitude to the observed *Bcd* degradation rates (*k ∼* 10^*−*4^*s*^*−*1^) in the system, and after substituting the values of concentration *C* and receptor size *a* from ref. [44] into above equation

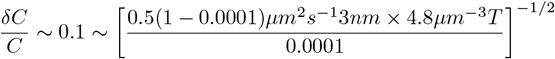

we see that the decoding of positional information from the *Bcd* morphogen gradient by its primary target gene *hb*, with the observed precision of ∼ 10 % takes *T ∼* 0.7 s, which is less than a second. From this one can readily calculate the on-rate of the *Bcd* morphogen for its target loci,

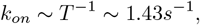

which is a very moderate on-rate. This is significant considering that lattice light sheet microscopy experiments have shown *Bcd* binding to chromatin sites in a highly transient manner, with specific binding events lasting on the order of a second, in all portions of the embryo with a rapid off rate such that its average occupancy at target loci is on-rate dependent [40].

